# Bovine mammary gland development: new insights into the epithelial hierarchy

**DOI:** 10.1101/251637

**Authors:** Laurence Finot, Eric Chanat, Frederic Dessauge

## Abstract

Milk production is highly dependent on the extensive development of the mammary epithelium, which occurs during puberty. It is therefore essential to distinguish the epithelial cells committed to development during this key stage from the related epithelial hierarchy. Using cell phenotyping and sorting, we highlighted three sub-populations that we assume to be progenitors. The CD49_f_^high^CD24^neg^ cells expressing *KRT14*, *vimentin* and *PROCR* corresponded to basal progenitors whereas the CD49_f_^low^CD24^neg^ cells expressing luminal KRT, progesterone and prolactin receptors, were of luminal lineage. The CD49_f_^low^CD24^pos^ cells had features of a dual lineage, with luminal and basal characteristics (CD10, ALDH1 and *KRT7* expression) and were considered to be early common (bipotent) progenitors. The mammary stem cell (MaSC) fraction was recovered in a fourth sub-population of CD49_f_^high^CD24^pos^ cells that expressed CD10/*KRT14* and *KRT7*. The differential ALDH1 activities observed within the MaSC fraction allowed to discriminate between two states: quiescent MaSCs and lineage-restricted “activated” MaSCs. The in-depth characterization of these epithelial sub-populations provides new insights into the epithelial cell hierarchy in the bovine mammary gland and suggests a common developmental hierarchy in mammals.

## INTRODUCTION

The mammary gland undergoes dynamic morphological changes over the lifetime of female mammals. At birth, bovine mammary parenchyma consists of a rudimentary duct network connected to a small cisternal cavity. At the onset of puberty, the mammary rudiment develops and starts to expand into the stroma upon stimulation by the ovarian steroid hormones, including estradiol and progesterone, and by growth factors (Yart et al., 2014). Ductal elongation occurs through the growth, development, and subsequent extension of terminal ductal lobular units (TDLU) in a process referred to as branching morphogenesis. In contrast to the long, infrequently branched ducts and terminal end buds found in the mammary gland of virgin mice, the mammary parenchyma of bovines develops into a compact, highly arborescent, parenchymal mass surrounded by a dense matrix of connective tissue (Akers, 2017). Bovine mammary TDLUs initially consist of solid cords of epithelial cells that penetrate into the stroma. As these cords extend into the mammary fat pad, lateral outgrowths emerge. This parenchymal development continues through puberty, until the mammary fat pad becomes filled. In addition, during gestation, the tissue continues its differentiation with the formation of lobulo-alveolar structures and the maturation of TDLUs in response to circulating hormones, notably prolactin. At the end of its development, the mammary epithelium has the appearance of an elaborate tree of ducts and alveoli. After parturition, the alveolar epithelium starts to be fully functional, with mammary epithelial cells secreting milk proteins into the lumen of the alveoli for lactation (McBryan and Howlin, 2017).

The ability of the mammary gland to undergo many cycles of lactation, with their stages of tissue proliferation and involution, suggests that the epithelial compartment contains resident cells capable of generating the entire epithelial architecture. Evidence for the existence of mammary stem cells (MaSCs) has been primarily derived from transplantation studies with murine mammary tissues. These studies revealed that the ductal architecture could be regenerated *in vivo* when isolated parenchymal explants were transplanted into cleared mammary fat pads (Deome et al., 1959; Ormerod and Rudland, 1986; Smith and Medina, 1988). More recent assays showed that an entire and functional mammary gland can be reconstituted from the transplantation of the progeny of a single “stem-like” cell (Shackleton et al., 2006; Stingl et al., 2006). Since these pioneering demonstrations, many studies in murine and human species have focused on identifying and isolating MaSC populations in order to establish the hierarchical cell organization and the molecular players in the regulation of the epithelium (Visvader and Stingl, 2014; Dontu and Ince, 2015). The epithelial hierarchy can be described as a pyramidal setup of the epithelial cell populations with stem cells at the apex and differentiated mature cells at the base of the pyramid. Between these two cell populations are the multiple progenitors that originate from the division and activation of stem cells and that progressively differentiate into mature cell lineages. Of note, the mammary structures are described as being composed of two major lineages: the luminal and basal cells, the latter including the myoepithelial cells. Luminal and basal cells can be distinguished by either their location in the epithelial structure or their protein expression profiles. Cells of these two lineages are considered immature during development as compared to the differentiated (mature) cells that constitute the functional secretory tissue.

In contrast, in bovines, only a few groups have attempted to elucidate the epithelial hierarchy *via* the identification of progenitor/stem cell populations (Martignani et al., 2009; Rauner and Barash, 2012). We recently participated in this research effort by providing original data on the mammary epithelial hierarchy committed to lactation during a lactation cycle in bovines (Perruchot et al., 2016). In this study, we used flow cytometry analysis and fluorescence activated cell sorting based on the expression of classic markers previously identified in the murine, human and bovine species. These markers are cell surface proteins, including the cluster of differentiation (CD) 24 (heat-stable antigen), CD29 (ß1-integrin) or CD49f (α6-integrin), and CD10 (Sleeman et al., 2006; Inman et al., 2015). These approaches led us to isolate putative populations of MaSCs, a prerequisite for further study of these target cell populations.

Research on MaSC biology in dairy mammals is important and relates to their potential use to improve animal robustness through the enhancement of lactation efficiency and infection resistance. In bovines, appropriate expansion and regulation of MaSCs may benefit mammogenesis, milk yield and tissue regenerative potential, making animals more robust (Capuco et al., 2012). A better understanding of the epithelial hierarchy at each developmental stage is therefore a prerequisite for the optimization of lactation in cows. Until now, literature describing the epithelial cell populations at key developmental stages (after puberty) and the regulators governing the bovine epithelial hierarchy has been scant. In this context, our study aims to further characterize the cells that make up the epithelial lineage at the branching morphogenesis stage in bovines, in order to provide new insights into the epithelial hierarchy.

## RESULTS

### Discrimination between cell sub-populations within the mammary epithelium of pubertal cows using the cell surface markers CD49_f_, CD24 and CD10

As puberty is a key period of mammary gland development, during which the different epithelial lineages, basal/myoepithelial and luminal cells, are committed to the process of branching morphogenesis and are identifiable, we used mammary gland samples from pubertal cows for our study.

In agreement with this, tissue staining with hematoxylin and eosin showed numerous neo-formed ductal and alveolar structures constituting an epithelium that largely formed the mammary parenchyma (Fig. S1). To identify the cell sub-populations of the epithelial lineages acting in the building of this parenchyma in the most exhaustive way possible, we focused our analysis on three cell surface markers that are well known to be specific for mammary epithelial cells: CD49_f_, CD24 and CD10. To validate our approach, we first analyzed the *in situ* localization of the cells expressing these markers by immunofluorescence. As shown in Figure 1, cells of the ductal trees at the origin of future TDLUs were clearly stained by anti-CD49_f_ antibodies (Fig.1, left panels). The outer cells of these epithelial structures formed a monolayer and were strongly stained at their basal side, whereas the inner cells were weakly stained. In contrast, cells expressing CD24 were scarce and scattered throughout the tissue slice (Fig.1, middle panels). Indeed, some cells were clearly found within the stromal tissue while others were localized in or near the lumen of the ducts, or close to the outer cell layer. As for CD10, which has been described as a cell surface marker of basal cells, it was clearly expressed by cells surrounding the developing duct structures (Fig.1, right panels). In this case, stained cells were exclusively localized to the outer epithelium layer, or sometimes appeared in small clusters (see the little structure at the top right of the image (Fig.1, right panels). These immuno-histological results having confirmed the relevance of using these markers, we decided to evaluate the proportion of each cell sub-population of the mammary tissue expressing them by flow cytometry.

**Fig. 1.**
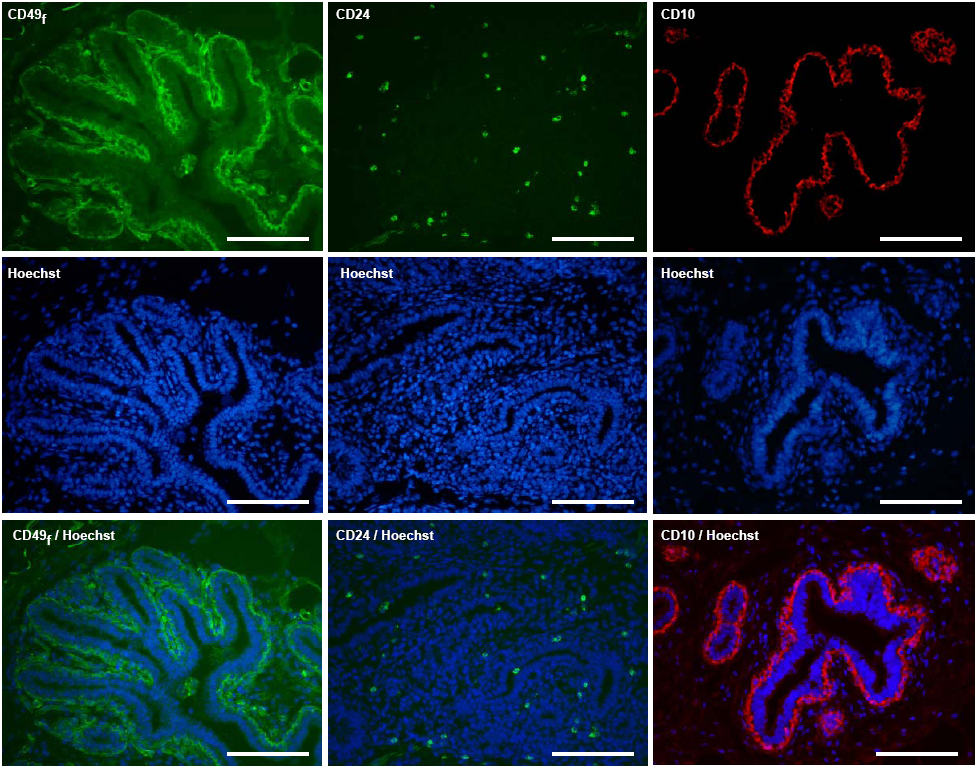
The cell surface markers CD49_f_, CD24 and CD10 are located in the luminal and basal cells within the ductal mammary epithelium of cows at puberty. Cryo‐ (CD49_f_ and CD24) and paraffin sections (CD10) from the mammary tissue of pubertal cows were processed for immunofluorescence for the indicated antigens. Nuclei were counterstained with Hoechst 33342. The basal membrane of the outer cell layer of the epithelium was highly stained for CD49_f_ whereas luminal cells were weakly stained (left panels, green). CD24-positive cells were scattered within the mammary epithelial structures (middle panels, green). Antibodies against CD10 nicely stained the outer cells of the developing ductal structures (right panels, red). Images are representative of three cows. Scale bars, 100μm.

As shown in the cytometric profile of CD49_f_ expression (Fig.2, upper plot), 62% (± 1.8%) of total single cells prepared from the mammary tissue of pubertal cows were CD49_f_^pos^ cells. Moreover, it was possible to distinguish two distinct sub-populations within these cells: the CD49_f_^low^ (42.2%) and CD49_f_^high^ (25%) sub-populations. To further identify the cell types that compose the mammary gland tissue of the pubertal cow, total single cells were sorted based on CD49_f_ expression. A set of proteins known to be specifically expressed in the epithelial lineage was then quantified in both negative and positive cell sub-populations by western blotting. What was first noticeable was the higher expression level of all epithelial lineage protein markers in the CD49_f_^pos^ cells compared to the CD49_f_^neg^ cells (fig.3A and 3B). First note that only the CD49_f_^pos^ cells expressed the epithelial cadherin (CDH1, Fig.3A, left graph), a protein involved in epithelial cell-to-cell adhesion. Moreover, these cells significantly overexpressed the basal marker CD10 when compared to the CD49_f_^neg^ cells (Fig. 3B). Similarly, the basal marker keratin (KRT) 14 and the myoepithelial marker alpha-smooth muscle actin (αSMA) were both absent in CD49_f_^neg^ cells, while these proteins were found in substantial amounts in CD49_f_^pos^ cells. Interestingly enough, we also observed that only the cells of the CD49_f_^pos^ sub-population expressed the luminal KRT7, KRT19 and KRT18 (see also Fig. S2 for the *in situ* lineage-specific localization of KRT). Altogether, these data strongly suggested that CD49_f_ cell sorting at least allowed the recovery of epithelial cells of both basal and luminal origins.

**Fig. 2.**
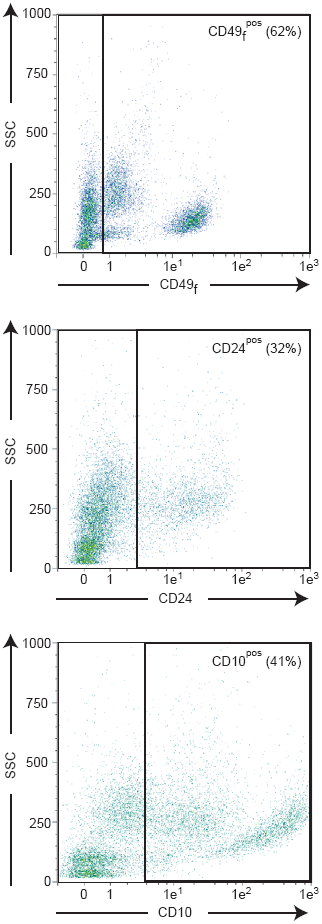
Distinct CD49_f_, CD24 or CD10 expression characterize sub-populations of bovine mammary epithelial cells. Dissociated cells from the mammary tissue of pubertal cows were stained with either anti-CD49_f_–FITC (CD49_f_), anti-CD24-APC (CD24) or anti-CD10-PE Vio770 (CD10) antibodies and analyzed by flow cytometry. Each gating shows the positive cells. The mean percentage of positive cells determined from the flow cytometric profiles of three independent experiments (3 cows) is indicated. Abbreviation: SSC, Side Scatter light.

When cells were analyzed for CD24 expression, a unique heterogeneous population of CD24^pos^ cells was observed (Fig.2, middle plot). It accounted for 32% (± 9.8%) of total single mammary cells. Western blotting showed that the epithelial marker CDH1 was expressed in both CD24^neg^ and CD24^pos^ cells (fig.3B) but was much more abundant in the latter cells. As a whole, the CD24^neg^ cells preferentially expressed the basal markers, i.e., CD10, αSMA and KRT14, whereas the luminal markers were more highly expressed in the CD24^pos^ cells. Indeed, both CD24^neg^ and CD24^pos^ cells expressed KRT7, KRT18 and KRT19, but all the luminal keratins were expressed at significantly higher levels in the CD24^pos^ population (Fig. 3A, middle graph and Fig.3B). We concluded that CD24 is a marker that allows the distinction of epithelial sub-populations within the basal and the luminal lineage.

**Fig. 3.**
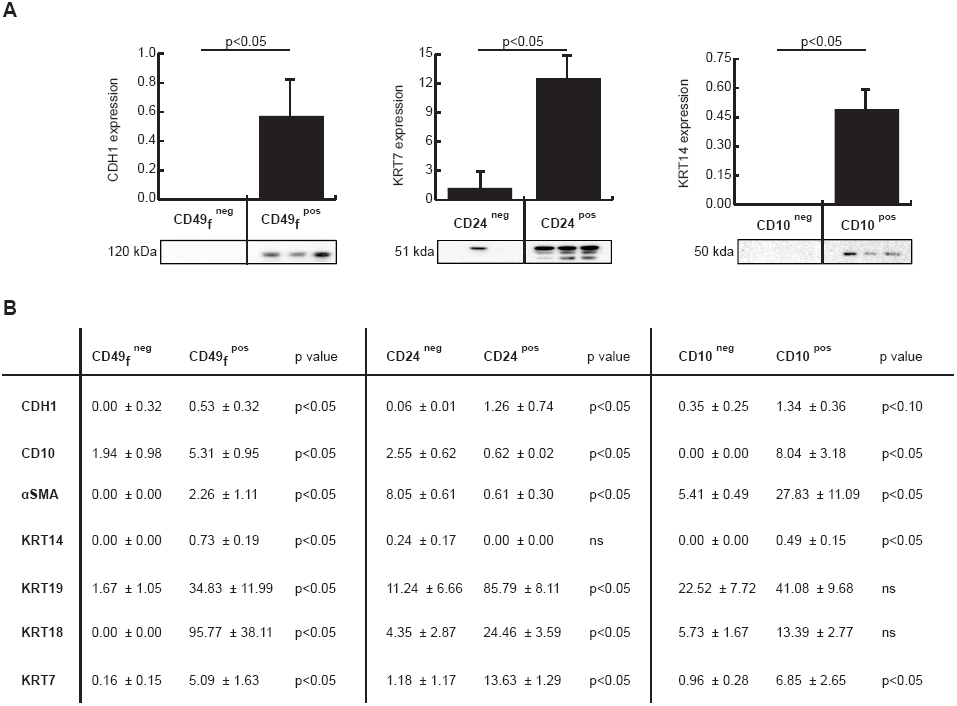
The expression of CD49_f_, CD24 and CD10 correlates to epithelial cell lineages of the bovine mammary gland. Sub-populations were sorted from the mammary tissue of pubertal cows according to the level of expression of either CD49_f_, CD24 or CD10 and total protein extracts were analyzed by western blotting with the indicated antibodies. The ECL signal was quantitated and the amount of each protein was expressed as percent of total proteins. Three independent experiments were performed (3 cows) and data are presented as means ± SEM. (**A**) Markers of the epithelial cell lineage distinguish the sorted cell sub-populations. The epithelial cadherin protein CDH1 was only present in the CD49_f_^pos^ cells, while the luminal marker protein KRT7 and the basal marker protein KRT14 were expressed in the CD24^pos^ and CD10^pos^ sub-populations, respectively. Relative molecular masses (kDa) are indicated. (**B**) Table summarizing western blotting data for protein markers of the epithelial cell lineages. Cells dissociated from mammary tissue of pubertal cows were stained with anti-CD49f, anti-CD24 and anti-CD10 antibodies. Positive and negative cell populations were collected by cell sorting and proteins were extracted to perform Western blotting. The level of expression of the indicated proteins was quantified and expressed as the percentage of total loaded protein ±SEM. Statistical analysis was performed using the Mann-Whitney U test. P value indicates significant differences (p<0.05), trends (p<0.1) and non-significant (ns) differences. Abbreviations: CDH1, E-cadherin; αSMA, alpha Smooth Muscle Actin; KRT14, Keratin 14; KRT19, Keratin 19; KRT18, Keratin 18; KRT7, Keratin 7.

Finally, when analyzing the cells for CD10 expression (Fig.2, bottom plot) we identified within the CD10^pos^ cells, two cell sub-populations expressing either low (CD10^low^) or high levels of CD10 (CD10^high^), the sum of which accounted for 41% (± 7.7%) of total mammary cells. Following cell sorting, we found that KRT14 was only present in the CD10^pos^ cells (Fig.3A, right graph and Fig.3B). In addition, αSMA was almost 6-fold more abundant in the CD10^pos^ population than in the CD10^neg^ population (27.8% ± 11% *vs* 5.4% ± 0.5%). Interestingly, the luminal KRT19, KRT18 and KRT7 were expressed in both the CD10^neg^ and CD10^pos^ cell sub-populations with no significant difference, except for KRT7, which was expressed at 6-fold higher level in the CD10^pos^ cells than in the CD10^neg^ cells (6.85% ± 2.6% *vs*. 0,96% ± 0.2%). The keratins seemed differentially expressed in luminal cells and may be most likely expressed according to their differentiation status. In summary, our data confirm that CD10 expression is characteristic of basal cells, making it a pertinent marker to discriminate the basal lineage from the luminal lineage.

### Determination of the cell sub-populations involved in mammary gland development at puberty

To further delineate the different cell sub-populations involved in the development of the mammary gland in pubertal cows, we analyzed all combinations of cell co-staining with CD49_f_, CD24 and CD10 by flow cytometry. Co-staining for CD49_f_ and CD24 revealed four distinct positive cell sub-populations in addition to the double-negative population (Fig.4, upper plot). The major cell sub-population was CD49_f_^pos^CD24^neg^ (42% ± 0.8% of total cells). These cells, however, were equally distributed in two sub-populations according to their fluorescence intensity, the CD49_f_^low^ (21.3% ± 0.8% of total cells) and CD49_f_^high^ cells (21.1% ± 2.3% of total cells, Table 1). The CD49_f_^pos^CD24^pos^ sub-populations represented 20% (± 3.7%) of total single cells with a large proportion of CD49_f_^low^CD24^pos^ cells (Fig.4, upper plot and Table 1). Interestingly, each of these sub-populations (CD49_f_^low^CD24^pos^, CD49_f_^low^CD24^neg^ and CD49_f_^high^CD24^neg^) approximately accounted for one third of the total CD49_f_^pos^ cells (see Fig. S4). Finally, we found that only 2% (± 0.1) of total single cells were CD49_f_^neg^CD24^pos^. Co-staining for CD49_f_ and CD10 revealed five distinct sub-populations (Fig.4, middle plot). Double-negative cells accounted for 23.4% (± 3.8%) of total single cells, 14.2% (± 4.4%) were CD49_f_^neg^CD10^pos^ and 36.5% (± 2.5%) were double-positive. Within the CD49_f_^pos^ populations, several sub-populations were well identifiable by their expression of both CD10 and CD49_f_ (13.7% (± 1.4%) of CD49_f_^low^CD10^pos/low^ and 17% (± 3.9%) of CD49_f_^high^CD10^pos/high^, see Table 1). Finally, co-staining for CD10 and CD24 (Fig.4, bottom plot) revealed heterogeneous sub-populations (Table 1). Altogether, these data highlighted the multiple cell sub-populations present within the mammary tissue during pubertal development.

**Fig. 4.**
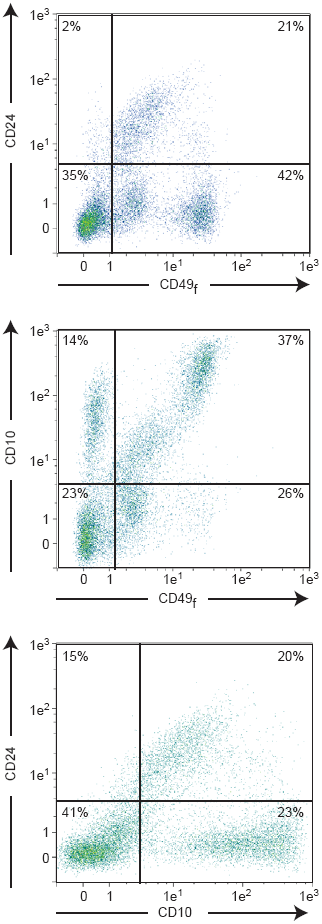
Sub-populations of epithelial cells cohabitate within the developing bovine mammary epithelium. Dissociated cells from the mammary tissue of pubertal cows were co-stained with either anti-CD49_f_-FITC and anti-CD24-APC antibodies, anti-CD49_f_-FITC and anti-CD10-PE Vio770 antibodies, or anti-CD10-PE Vio770 and anti-CD24-APC antibodies, and analyzed by flow cytometry. Each gating shows the positive cells; positive cells are located to the right of the gating on the x-axis and above the gating on the y-axis. Sub-populations of epithelial cells were distinguished according to the intensity of the cell surface marker expression (low vs. high). The mean percentage of cells in each quadrant (percentage of total cells) determined from the flow cytometric profiles of three independent experiments (3 cows) is indicated.

**Table 1.** Results of flow cytometry analysis for CD49f, CD24 and CD10 expression in mammary gland of heifers

| Monostained Populations | % ± SEM |
| --- | --- |
| **CD49<sub>f</sub> Populations** |  |
| CD49<sub>f</sub><sup>neg</sup> | 37.4 ± 1.8 |
| CD49<sub>f</sub><sup>pos</sup> | 62.6 ± 1.8 |
| **CD24 Populations** |  |
| CD24<sup>neg</sup> | 67.4 ± 9.2 |
| CD24<sup>pos</sup> | 32.5 ± 9.8 |
| **CD10 Populations** |  |
| CD10<sup>neg</sup> | 62.9 ± 13.7 |
| CD10<sup>pos</sup> | 41.3 ± 7.7 |
| **Doublestained Populations and Subpopulations** |  |
| **CD49<sub>f</sub> /CD24 Populations** |  |
| CD49<sub>f</sub><sup>neg</sup> CD24<sup>neg</sup> | 35.3 ± 1.7 |
| CD49<sub>f</sub><sup>neg</sup> CD24<sup>pos</sup> | 2.2 ± 0.1 |
| CD49<sub>f</sub><sup>pos</sup> CD24<sup>neg</sup> | 41.9 ± 2.7 |
| \| CD49<sub>f</sub><sup>low</sup> CD24<sup>neg</sup> | 20.8 ± 0.8 |
| \| CD49<sub>f</sub><sup>high</sup> CD24<sup>neg</sup> | 20.8 ± 2.3 |
| CD49<sub>f</sub><sup>pos</sup> CD24<sup>pos</sup> | 20.6 ± 3.7 |
| \| CD49<sub>f</sub><sup>low</sup> CD24<sup>pos</sup> | 16.8 ± 3.2 |
| \| CD49<sub>f</sub><sup>high</sup> CD24<sup>pos</sup> | 3.4 ± 0.4 |
| **CD49<sub>f</sub> /CD10 Populations** |  |
| CD49<sub>f</sub><sup>neg</sup> CD10<sup>neg</sup> | 23.4 ± 3.8 |
| CD49<sub>f</sub><sup>neg</sup> CD10<sup>pos</sup> | 14.2 ± 4.4 |
| CD49<sub>f</sub><sup>pos</sup> CD10<sup>neg</sup> | 25.8 ± 3.7 |
| \| CD49<sub>f</sub><sup>low</sup> CD10<sup>neg</sup> | 20.7 ± 3.2 |
| \| CD49<sub>f</sub><sup>high</sup> CD10<sup>neg</sup> | 2.1 ± 0.3 |
| CD49<sub>f</sub><sup>pos</sup> CD10<sup>pos</sup> | 36.5 ± 2.5 |
| \| CD49<sub>f</sub><sup>low</sup> CD10<sup>pos</sup> | 13.7 ± 1.4 |
| \| CD49<sub>f</sub><sup>high</sup> CD10<sup>pos</sup> | 17.1 ± 3.9 |
| **CD10 /CD24 populations** |  |
| CD10<sup>neg</sup> CD24<sup>neg</sup> | 40.9 ± 1.9 |
| CD10<sup>neg</sup> CD24<sup>pos</sup> | 15.4 ± 1.8 |
| CD10<sup>pos</sup> CD24<sup>neg</sup> | 23 ± 6.2 |
| CD10<sup>pos</sup> CD24<sup>pos</sup> | 20 ± 6.4 |

Data of cellular populations and sub-populations are expressed as the mean percentage of cells ± SEM from three independent experiments (3 heifers)

### Characterization of the cell sub-populations composing the mammary epithelial hierarchy

As mammary stem cells and progenitors were reported to belong to a subset of CD49_f_^pos^CD24^pos^ cells, we decided to further depict the CD49_f_ and CD24 sub-populations by further investigating their phenotype. These sub-populations were analyzed for both CD10 expression and aldehyde dehydrogenase 1 (ALDH1) activity by flow cytometry (Fig.5). We found that the CD49_f_^low^CD24^neg^ cells were predominantly negative for CD10 (Fig.5 middle, left plot) whereas almost all CD49_f_^high^ cells expressed CD10 (Fig.5 middle, right plots). Within the CD49_f_^low^CD24^pos^ sub-population, 75% of the cells were positive for CD10 (Fig.5 middle, second left plot). Interestingly enough, a correlation was observed between the intensity of CD49_f_ and CD10 fluorescence, all CD49_f_^high^ cells being CD10^high^. Similarly, we evaluated the activity of ALDH1 in the aforementioned CD49_f_^pos^ sub-populations. Indeed, ALDH1 activity has been previously identified as a marker of luminal cells and it has been shown to distinguish progenitor from mature mammary luminal cells in some species (Eirew et al., 2012). We found that 70 to 86% of the CD49_f_^low^ cells, namely the CD49_f_^low^CD24^neg^ and the CD49_f_^low^CD24^pos^ cells, exhibited ALDH1 activity, as well as 70 % of the CD49_f_^high^ CD24^pos^ cells. It is therefore reasonable to assume that these three sub-populations belong or are related to the luminal lineage.

**Fig. 5.**
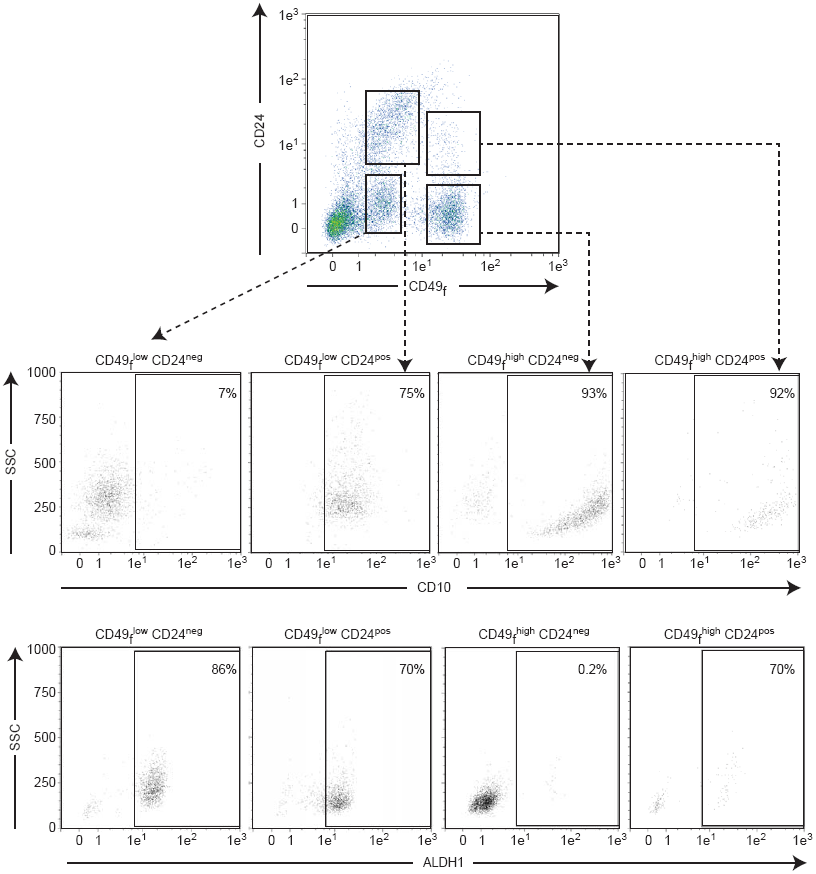
Sub-populations of epithelial cells exhibit distinct lineage types in the developing bovine mammary gland. Cells dissociated from pubertal bovine mammary tissue were co-stained with anti-CD49_f_ ‐FITC (CD49_f_) and anti-CD24-APC (CD24) antibodies and analyzed by flow cytometry (upper plot). Cells expressing low or high intensities of CD49_f_ and/or CD24 were sorted and subjected to FACS analysis for either CD10 expression (middle plots) or ALDH1 activity (lower plots). Representative flow cytometry analysis plots for CD10 or ALDH1 expression for each sub-population are shown. Gating on quadrants highlight positive cells and the mean percentage of cells in each quadrant (percentage of total cells) determined from the flow cytometric profiles of three independent experiments (3 cows) is indicated. Abbreviations: SSC, Side Scatter Light; ALDH1, Aldehyde dehydrogenase.

We next investigated the expression of target genes by RT-qPCR. Those included the keratins, vimentin, some stem cell markers picked from the literature, and hormonal receptor genes as indicators of differentiation (Table 2). Hormone receptivity of the mammary tissue was assessed beforehand by immunofluorescence staining of the progesterone (PR) and estradiol (ERα) receptors (see Fig. S3A). This revealed their presence in the epithelial cells and therefore the sensitivity of these cells to hormones (22% ± 2.4% and 11% ± 1%, for PR and ERα-stained cells, respectively) (Fig. S3B). We also found that the genes known to be expressed by stromal cells, namely *vimentin, ALDH1* and the *Protein C receptor* (*PROCR*) were expressed significantly more expressed in the CD49_f_^neg^CD24^neg^ sub-population than in the other sub-populations. Additionally, this sub-population under-expressed genes of the KRT family and the differentiation/receptivity markers compared to the other sub-populations. On the other hand, significant differences in gene expression were found between the CD49_f_^pos^ sub-populations. Indeed, the two CD49_f_^low^ sub-populations expressed higher levels of *KRT19, KRT18 and KRT7* compared to the CD49_f_^neg^ sub-population, confirming their luminal origin. However, the CD49_f_^low^CD24^neg^ and CD49_f_^low^CD24^pos^ sub-populations composing the CD49_f_^low^ populations presented differences in KRT expression (2- and 2.6-fold more abundant for *KRT19* and *KRT18*, respectively, in the CD49_f_^low^CD24^neg^ sub-population than in the CD49_f_^low^CD24^pos^ sub-population) and in to their hormonal receptivity (2.5-fold more abundant for *PR* and *prolactin receptor* (*PRLR)* in the CD49_f_^low^CD24^neg^ sub-population than in the CD49_f_^low^CD24^pos^ sub-population). The CD49_f_^low^CD24^neg^ sub-population was characterized by expression of the three luminal keratins and of both *PR* and *PRLR*. The CD49_f_^low^CD24^pos^ sub-population especially expressed the luminal *KRT7*, the stemness markers *ALDH1* and the receptivity markers *PR* and *E74-like factor 5 (ELF5)*. As for the CD49_f_^high^ sub-populations, they significantly expressed *KRT14*, confirming their basal origin. Finally, the CD49_f_^high^CD24^neg^ sub-population was characterized by a moderate abundance of the *vimentin* and *PROCR* genes whereas the CD49_f_^high^CD24^pos^ sub-population expressed the *KRT7, ALDH1 and ELF5* genes. In conclusion, each CD49_f_ CD24 sub-population exhibited a specific phenotype and molecular signature which allowed them to be catalogued in a lineage type.

**Table 2.** Gene expression levels in epithelial sub-populations.

|  | CD49<sup>f</sup><sup>neg</sup> CD24<sup>neg</sup> | CD49<sup>f</sup><sup>low</sup> CD24<sup>neg</sup> | CD49<sup>f</sup><sup>low</sup> CD24<sup>pos</sup> | CD49<sup>f</sup><sup>hi</sup> CD24<sup>neg</sup> | CD49<sup>f</sup><sup>hi</sup> CD24<sup>pos</sup> | p value |
| --- | --- | --- | --- | --- | --- | --- |
| **Cellular type markers** |  |  |  |  |  |  |
| *KRT14* | 0.014<sup>b</sup> | 0.034<sup>b</sup> | 0.149<sup>b</sup> | 4.060<sup>a</sup> | 4.046<sup>a</sup> | p<0.05 |
| *KRT19* | 0.001<sup>b</sup> | 0.064<sup>a</sup> | 0.035<sup>ab</sup> | 0.003<sup>b</sup> | 0.032<sup>ab</sup> | p<0.01 |
| *KRT18* | 0.002<sup>b</sup> | 0.243<sup>a</sup> | 0.095<sup>b</sup> | 0.022<sup>b</sup> | 0.080<sup>b</sup> | p<0.01 |
| *KRT7* | 0.000<sup>b</sup> | 0.032<sup>a</sup> | 0.028<sup>a</sup> | 0.002<sup>b</sup> | 0.030<sup>a</sup> | p<0.01 |
| *Vimentin* | 1.437<sup>a</sup> | 0.085<sup>cd</sup> | 0.030<sup>d</sup> | 0.944<sup>b</sup> | 0.490<sup>c</sup> | p<0.01 |
| **Stemness markers** |  |  |  |  |  |  |
| *NOTCH1* | 0.010<sup>a</sup> | 0.006<sup>a</sup> | 0.004<sup>a</sup> | 0.009<sup>a</sup> | 0.013<sup>a</sup> | ns |
| *ALDH1* | 0.024<sup>a</sup> | 0.003<sup>bc</sup> | 0.008<sup>b</sup> | 0.000<sup>c</sup> | 0.008<sup>b</sup> | p<0.01 |
| *PROCR* | 0.0034<sup>a</sup> | 0.000<sup>c</sup> | 0.000<sup>c</sup> | 0.0005<sup>b</sup> | 0.0001<sup>c</sup> | p<0.01 |
| **Differentiation / Receptivity markers** |  |  |  |  |  |  |
| *Estrogen Receptor* | 0.009<sup>a</sup> | 0.082<sup>a</sup> | 0.039<sup>a</sup> | 0.006<sup>a</sup> | 0.033<sup>a</sup> | ns |
| *Progesterone Receptor* | 0.000<sup>b</sup> | 0.041<sup>a</sup> | 0.017<sup>ab</sup> | 0.002<sup>b</sup> | 0.010<sup>b</sup> | p<0.01 |
| *Prolactin Receptor* | 0.003<sup>c</sup> | 0.399<sup>a</sup> | 0.160<sup>b</sup> | 0.023<sup>c</sup> | 0.155<sup>b</sup> | p<0.01 |
| *ELF5* | 0.001<sup>b</sup> | 0.003<sup>b</sup> | 0.041<sup>a</sup> | 0.001<sup>b</sup> | 0.061<sup>a</sup> | p<0.01 |

Cells dissociated from heifer mammary tissue were co-stained with anti-CD49f-FITC and anti-CD24-APC antibodies and sorted based on the level of markers expression (low and high). The level of expression of the indicated genes was measured by RT-qPCR and normalized to the amount of *RPLP0* transcript, the most stable gene in a panel of 3 reference genes. Data are expressed as the mean of Delta Delta Ct calculation. Different letters (a-d) indicate significant differences.

## DISCUSSION

After puberty, each estrous cycle is accompanied by periods of enhanced cell proliferation and differentiation in the mammary gland until the fat pad is filled with parenchymal tissue. However, we believe that the post-pubertal stage is a much wiser period in which to able to identify the most epithelial cell categories, including progenitor cells. This is of substantial importance as the branching process during puberty evolves and because the phenotype of the epithelial sub-populations involved at the beginning of puberty may well change during the progression of the branching process. That is why we deliberately chose to work on pubertal animals. Analysis of the expression by bovine mammary epithelial cells at puberty of the specific cell surface markers CD49_f,_ CD24 and CD10 using flow cytometry allowed the identification and isolation of prospective key cell sub-populations. Of course, it was of the utmost interest to further analyze the molecular signatures of these sub-populations to improve our knowledge of the bovine mammary epithelial cell hierarchy.

### The majority of the epithelial cells committed to mammary development at puberty are progenitors

We first found that the CD49_f_^high^CD24^neg^ sub-population expressed KRT14, a well-known marker of the basal lineage classically associated with myoepithelial cells (Dairkee et al., 1988; Safayi et al., 2012). This sub-population also substantially expressed CD10, another marker of basal cells (Safayi et al., 2012). Finally, immunohistological observation revealed that the cells of the outer epithelium layer were strongly stained at their basal side by anti-CD49_f_ antibodies. In summary, these data indicate that the CD49_f_^high^CD24^neg^ sub-population is from the basal lineage. This is in agreement with a previous bovine study on the characterization of the epithelial cells present in the mammary gland a few months after birth (Rauner and Barash, 2012). More recently, this group reported that, at early developmental stages, the basal cells were CD49_f_^pos^CD24^neg^, and specified that their phenotype was CD49_f_^high^CD24^neg^ (Rauner and Barash, 2016). In the present study, we further characterized this basal CD49_f_^high^CD24^neg^ sub-population by, notably, studying the expression of the *vimentin* and *PROCR* genes. Indeed, vimentin filaments are expressed, *inter alia*, in the basal epithelial cell population of the mammary gland (Peuhu et al., 2017) and it has recently been demonstrated that *vimentin* deficiency in *vimentin* KO mice affects mammary ductal development by altering progenitor cell activity (Peuhu et al., 2017). This suggested a regulatory role of vimentin in the basal MaSC/progenitor cell population. Here, the observation that the CD49_f_^high^CD24^neg^ cells expressed *vimentin* prompted us to propose that this sub-population is probably progenitor cells. This is also supported by the observation that these cells expressed high levels of *PROCR*. Indeed, although PROCR was originally studied as a stem cell marker in hematopoiesis, this protein was also found to be relatively abundant in the basal cells of murine mammary epithelium (Wang et al., 2015). In this latter study, PROCR was suggested to be a marker of mammary stem cells, a possibility that was previously envisioned in a model of human breast cancers, in which the receptor was one of the molecular markers for stem/progenitor-like populations (Shipitsin et al., 2007). Taking a middle-ground position, we can claim that the CD49_f_^high^CD24^neg^ cell sub-population accounts for the basal progenitor cells.

Immunofluorescence analysis showed that cells localized to the inner epithelium layer expressed low levels of CD49_f_. Also, our cytometric profiles showed two mammary epithelial cell sub-populations that expressed low levels of CD49_f_. This is in agreement with the aforementioned study in bovines (Rauner and Barash, 2016) and with studies in mouse, in which the luminal population was reported as being CD49_f_^low^ (Asselin-Labat et al., 2006; Rauner and Barash, 2016; O’Leary et al., 2017). In addition to these data, we showed here by western blotting that KRT19, KRT18 and KRT7 were expressed by the CD49_f_^pos^ cells, including the CD49_f_^low^ cells. The abundance of these keratins was also demonstrated at the mRNA level in the two CD49_f_^low^ sub-populations (CD49_f_^low^CD24^neg^ and CD49_f_^low^CD24^pos^). In the mammary gland, the relative expression of specific keratins by distinct epithelial cells is well established and is cell lineage-specific. They are therefore classically used to distinguish the luminal cells from the basal cells. Indeed, luminal cells of the epithelium express KRT7, KRT8, KRT18 and KRT19 cells, whereas basal cells express KRT5 and KRT14. Taken together, our data confirm that the CD49_f_^low^ populations belong to the luminal lineage. On the other hand, we showed that the CD49_f_^low^ sub-populations can be distinguished by the expression of CD24. Furthermore, we found that both CD49_f_^low^CD24^neg^ and CD49_f_^low^CD24^pos^ sub-populations exhibited ALDH1 activity, a feature that identifies the differentiation status of the luminal cells. Indeed, a previous study in human mammary gland demonstrated that ALDH1 activity was upregulated at the transition of progenitor cells into the luminal lineage, making it possible to define the luminal progenitor cells (Eirew et al., 2012). ALDH1 activity has also been used in the bovine model to define luminal-restricted progenitors (Martignani et al., 2010) and in the mouse model to distinguish the relatively undifferentiated luminal progenitors from the differentiated ones (Shehata et al., 2012). Finally, we found that both CD49_f_^low^CD24^pos^ and CD49_f_^low^CD24^neg^ cells expressed high levels of *KRT7*, a marker of immature luminal cells (Lichtner et al., 1991). From these studies and our data, we conclude that the two CD49_f_^low^ sub-populations are luminal progenitors.

Of note, the CD49_f_^low^CD24^neg^ sub-population expressed high levels of the *PR* and *PRLR* genes. Many studies have reported that mammary development is triggered at puberty by the main steroid hormones estradiol and progesterone (for review see (McBryan and Howlin, 2017)). These hormones may act jointly or independently, suggesting a spatio-temporal regulation by each hormone. Indeed, experiments with *PR*-deficient mice demonstrated that, at puberty, progesterone is not essential for ductal elongation but is critical in inducing side-branching (Atwood et al., 2000). This observation suggests that progesterone, independent from estradiol, could intervene late in branching morphogenesis to promote side branching and then in the formation of lobulo-alveolar structures (Brisken and Ataca, 2015). Moreover, it has been found that a large number of luminal cells are PR-positive in adult virgin mice at an advanced stage of puberty (Seagroves et al., 2000). We made similar observations in the bovine mammary tissue of pubertal cows by immunofluorescence, with the PR staining being restricted to luminal cells. As to the key role of prolactin at this advanced stage of puberty, it has been found that deletion of the *PRLR* in mice resulted in defects in side branching and further alveolar formation, proving the role of prolactin in branching morphogenesis (Ormandy et al., 2003). Conversely, overexpression of prolactin in mice has been shown to increase lateral ductal budding and to increase epithelial progenitor sub-populations (O’Leary et al., 2017). Hence, our finding that the CD49_f_^low^CD24^neg^ sub-population expressed *PR* and *PRLR* plus ALDH1 activity strongly suggests that these cells are “mature progenitors” differentiated to promote side branching and/or alveogenesis.

As discussed above, the second luminal sub-population we found, namely the CD49_f_^low^CD24^pos^ cells, expressed mainly *KRT7* and exhibited ALDH1 activity, two features showing both their luminal lineage and a progenitor state. Surprisingly, the cytometric analysis revealed that these cells also expressed the basal cell marker CD10. In many human studies, it has been shown that some progenitor cells have the ability to produce both luminal colonies (expressing KRT8) and mixed luminal/basal colonies (expressing KRT8 and KRT14) when cultured *in vitro*, suggesting the existence of a bipotent cell population (Villadsen et al., 2007; Stingl, 2009). Therefore, we hypothesize that the cells forming the CD49_f_^low^CD24^pos^ sub-population undoubtedly have dual lineage features. In addition, we found that these cells expressed *ELF5* and *PR*, two genes well known to be expressed by the luminal lineage. Interestingly, these genes have recently also been associated with the regulation of progenitor/stem cells. Indeed, although the transcription factor ELF5 is known to orient the fate of luminal cells during alveogenesis (Oakes et al., 2008), *ELF5* deficiency was also shown to lead to the accumulation of luminal/basal (bipotent) cells and to increase the MaSC-enriched cell population. This latter observation confirmed the regulatory role of *ELF5* in the level of stem cells/progenitors (Chakrabarti et al., 2012). Finally, a consistent enrichment of the *PR* transcript was also observed in bipotent progenitors in the normal human breast (Hilton et al., 2012). In summary, the dual lineage features of the CD49_f_^low^CD24^pos^ cells (CD10^+^/*KRT7*^+^) plus the expression of the *PR* and *ELF5* genes in these cells prompted us to consider this population to be an early common progenitor characterized by bipotency. One can conclude that the three sub-populations discussed above, each of them representing 1/3 of the total number of epithelial cells, are progenitors that differ in their lineage (bipotent, luminal or basal lineage).

### Two sub-populations co-exist in the MaSC fraction

In many species, whether human, murine or bovine, the stem cell population, referred to as the MaSC population, has been described as being CD49_f_^high^CD24^pos^ (Borena et al., 2013; Visvader and Stingl, 2014; Rauner et al., 2017). In our study, this cell sub-population represented 5.5% of total epithelial cells or 3.8% of total mammary cells. This relatively small percentage was consistent with what is usually reported for the MaSC-enriched fraction in the literature (5% of total mammary cells in mice and 2.43% in post-pubertal bovines (Osinska et al., 2014). Recently, we showed that the proportion of CD49_f_^high^CD24^pos^ cells in the bovine lactating mammary gland range from 0.7% to 3.3% (Perruchot et al, 2016). In the present study, we found that this sub-population also expressed the two basal markers CD10 and *KRT14*. This was consistent with the observation that MaSCs appeared localized to the basal compartment in several studies, sharing characteristics with the surrounding basal cells (Bachelard-Cascales et al., 2010; Van Keymeulen et al., 2011; Prater et al., 2014). This is most likely in order to maintain both their anchorage and survival in this tissue compartment. As observed previously (Dontu and Ince, 2015) and confirmed here, the MaSCs contained in the CD49_f_^high^CD24^pos^ sub-population formed mammospheres when cultured for 7 days in the presence of matrigel (data not shown). The above considerations strongly suggest that the CD49_f_^high^CD24^pos^ sub-population we highlighted in the present work is the MaSC fraction. However, after in-depth analysis of the cytometry data, although these cells were homogeneous for CD10 expression, only 70% (corresponding to 3.8% of total epithelial cells) exhibited ALDH1 activity, whereas 30% (1.7% of total epithelial cells) had no ALDH1 activity. This suggests that the MaSC fraction contains two sub-populations, supporting the notion that stem cells are heterogeneous. This notion has recently been raised in an elegant study of the murine MaSCs (Scheele et al., 2017) in which the dynamics of branching morphogenesis were monitored by highlighting the behaviour of the different lineage-committed MaSCs using a “confetti” cell strategy. It emerged that MaSCs may be heterogeneous. Indeed, it was concluded that a pool of MaSCs is engaged in the development of the tissue whereas another stays quiescent. From this, we can hypothesize that the MaSC sub-populations exhibiting ALDH1 activity represent the lineage-restricted “activated” MaSC whereas the second sub-population probably contains the quiescent cells. If this is the case, the expression of *KRT7* and *ELF5* could also be attributed to the “activated” pool of MaSCs, which, with this commitment feature, could be at the origin of the bipotent cell population.

The data gathered in this study are consistent with those reported for earlier developmental stages of the bovine mammary gland (Rauner and Barash, 2012; Rauner and Barash, 2016). However, there are some differences, notably concerning the characteristics of sub-populations and the position of the bipotent cells in the hierarchy; we placed them between the MaSC sub-population and luminal progenitor cells. Of course, the different physiological stages of the animals used in the report mentioned above and in the present study, i.e., 7 months old (before puberty) *vs.* 17 months old (during puberty), might well explain the different phenotypic characteristics encountered for the various epithelial sub-populations.

### The epithelial cell hierarchy in the mammary gland at puberty

Based on our original results and according to the current literature, we conceived a mammary epithelial cell hierarchy scheme (Fig. 6). Of course, the stem cells, referred to as MaSCs and corresponding to the CD49_f_^high^CD24^pos^ sub-population, are placed at the top of this hierarchy. This MaSC pool is assumed to contain two sub-populations. The most undifferentiated cells (most likely the quiescent cells) are at the very top of the hierarchic tree. The second sub-population corresponds to the “activated-committed” MaSCs exhibiting early luminal markers (ALDH1, *KRT7*, *ELF5*) and basal markers (CD10 and *KRT14*). These cells are therefore close to bipotency. The “activated-committed” MaSCs generate the CD49_f_^low^CD24^pos^ cell sub-population, with phenotypic characteristics similar to those of the CD49_f_^high^CD24^pos^ cells. These are bipotent progenitor cells which have kept the expression of the same luminal markers as the “activated-committed” MaSC, and CD10 expression, but have lost *KRT14* expression. The comparison of the CD49_f_^low^CD24^pos^ and CD49_f_^low^CD24^neg^ cell sub-populations, with common expression of *KRT7* and *PR,* as well as ALDH1 activity, shows that these sub-populations are connected. Although the CD49_f_^low^CD24^neg^ cells have lost basal properties, they have acquired a panel of luminal keratins (*KRT19* and *KRT18*), clearly orienting them to a luminal fate. We speculated that the progressive differentiation of the bipotent cell sub-population into the luminal fate produces the luminal progenitor cells, corresponding to the CD49_f_^low^CD24^neg^ cell sub-population. The progressive differentiation of this sub-population, concretized here by the expression of the *PR* and *PRLR* receptors, makes these cells ready for the side branching process and/or alveolar formation. As to the basal/myoepithelial lineage, a distinct differentiation path may be involved. Indeed, the characteristics of the CD49_f_^high^CD24^neg^ cell sub-population are partly common to the CD49_f_^high^CD24^pos^ cell sub-population and are completely discordant with the others. These two cell sub-populations shared high expression of CD49_f_ and CD10, as well as expression of *KRT14*. Therefore, it is consistent to draw a basal lineage pathway in which the CD49_f_^high^CD24^pos^ cells (MaSC) supply the basal/myoepithelial progenitor cell sub-population (CD49_f_^high^CD24^neg^).

**Fig. 6.**
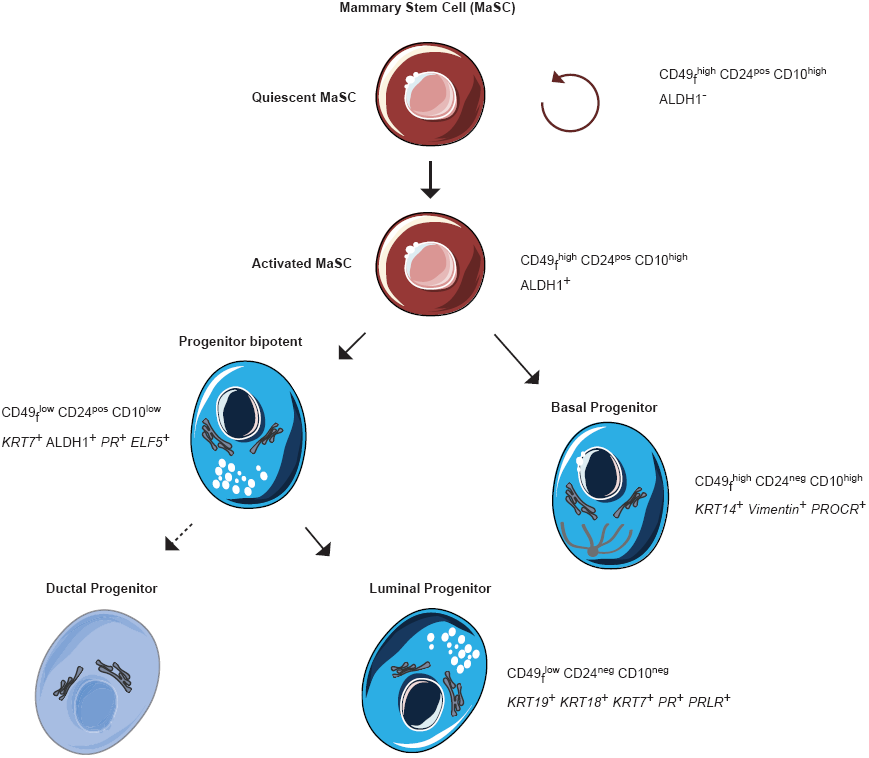
Schematic model of bovine mammary epithelial cell hierarchy.

It is confusing to compare mammary epithelial cell lineages between species from the literature, especially because investigators regularly use different cell markers (Stingl, 2009). Therefore, for the present study, we deliberately chose markers that have already been used in several species. In many schemas of mammary epithelial cell lineage proposed to date (mice, human, rat and other species), it is mentioned that stem cells shared the same characteristics (CD49_f_^high^CD24^pos^) (Asselin-Labat et al., 2008; Stingl, 2009; Rauner et al., 2017). Interestingly enough, the mammary epithelial cell hierarchy we propose here shares many common points with that proposed for the murine model (Visvader and Stingl, 2014). In addition, it is generally proposed that the stem cell population at the top of the hierarchy gives rise to a bipotent progenitor cell population and luminal or basal progenitors. It is therefore tempting to speculate from these studies and our data that the mammary epithelial cell hierarchy could be similar between mammals. Confirmation of these hypotheses as well as an evaluation of the epigenetic signature of each cell sub-population, supplemented by transplantation assays, could be relevant approaches to clarify the quiescent or activated status of each pool of MaSCs.

## MATERIALS AND METHODS

### Animals

The Holstein cows (*bos Taurus*) used in this study were housed at the experimental farm of Méjusseaume INRA-Rennes (France). The pubertal cows were sacrificed at 17 months of age at the slaughterhouse of Gallais Viande (Montauban-de-Bretagne, France) following standard commercial practices. The mammary glands were collected at the time of slaughter and immediately transported on ice to the laboratory to be sampled for further analyses.

### Mammary tissue sampling

Total parenchyma of the mammary gland was dissected and sampled. Samples destined for tissue dissociation were manually cut into small explants (≈1 mm^3^), suspended in 90% fetal bovine serum (10270-106; Gibco Invitrogen Saint Aubin, France)/ 10% dimethyl sulfoxide (DMSO, D2650, Sigma-Aldrich, Saint-Quentin Fallavier, France), and stored at -150°C. Tissue pieces (≈3 mm^3^) for RNA and protein extraction were snap frozen in liquid nitrogen and stored at -80 °C. For immunohistological analysis, tissue pieces (≈ 5 mm^3^) were fixed in 4% paraformaldehyde (FOR007OAF59001, VWR, Fontenay-sous-Bois, France) and were either mounted in OCT embedding compound (00411243, Labonord, Templemars, France) and frozen at -80°C, or dehydrated in ethanol and embedded in paraffin.

### Flow cytometry and cell sorting

Mammary tissue fragments were thawed and enzymatically dissociated as previously described (Perruchot et al., 2016) to obtain a single cell suspension. Dissociated cells were incubated with the relevant antibodies for 20 min at 4°C, washed and re-suspended in MACS buffer (130-091-222, Miltenyi Biotec, Paris, France) with 2% bovine serum albumin (130-091-376; Miltenyi Biotec) for flow cytometry analysis or cell sorting.

Flow cytometry was performed using a MACSQuant Analyzer 10 cytometer (Miltenyi Biotec). The controls and gating strategy used in the present study have been previously detailed (Perruchot et al, 2016). Note that isotype control antibodies were used as negative controls in the flow cytometry experiment. Data were analyzed using MACSQuantify analysis software (Miltenyi Biotec) and results expressed in percentage of cells out of 20,000 events.

ALDH1 activity was measured in 500.000 cells with the Aldefluor kit (01700, Stem cell technologies, Grenoble, France) according to the manufacturer’s recommendations. Cells were then centrifuged at 250G, re-suspended in Aldefluor assay buffer and labeled with antibodies against CD49_f_ and CD24.

For cell sorting, cells were incubated with the relevant antibodies for 20 min at 4°C in the dark. Single live cells were gated by DAPI exclusion and sorted on a BD FACS ARIA II flow cytometer (BIOSIT CytomeTRI technical Platform – Villejean Campus, Rennes, France). Sorted cells were centrifuged at 300G for 5 min at 4°C and stored at -80°C. The antibodies used are described in supplemental table S1.

### Protein extraction and Western Blotting

Proteins were extracted from sorted cell populations, quantified using the BCA assay kit (23227, Thermo Fisher, Illkirch, France) and analyzed by western blotting as previously described (Arevalo Turrubiarte et al., 2016), except that the amount of loaded protein was reduced to 2.5 μg. ECL signal was digitalized using the ImageQuant LAS4000 Imager digital system (GE Healthcare, Velizy-Villacoublay, France) and quantified with the ImageQuant TL software (GE Healthcare). An identical amount of each sample was analyzed in parallel by SDS-PAGE followed by Coomassie brilliant blue R-250 (161-0436, Biorad, France) staining. Gels were digitized and total protein in each track was quantified as described above for the ECL signal. ECL signals were expressed as the percentage of total protein. The antibodies used are described in supplemental table S1.

### mRNA extraction and quantitative PCR

RNA extraction was performed using the Nucleospin RNA XS kit (740902, Macherey-Nagel, Hoerdt, France) according to the manufacter’s instructions. Reverse transcription and quantitative PCR were performed as previously described (Perruchot et al, 2016). Raw cycle threshold (Ct) values obtained from StepOne Software version 2.3 (Applied Biosystems) were transformed into quantities using the delta delta Ct method. The endogenous control gene, the Ribosomal Protein Large P0 (*RPLP0*), was selected as the most stable gene within a panel of 3 genes (*18S rRNA, Ribosomal Protein S5* and *RPLP0*) using the Normfinder algorithm. The primers used in this study are described in supplemental table S2.

### Histological and immunohistochemical staining

Hematoxylin and eosin staining were performed on paraffin sections (8 μm) after rehydration as previously described (Perruchot et al., 2016). CD49_f_ and CD24 immunostaining (see below) were performed on frozen sections (5 μm) mounted on Superfrost Plus slides (4951PLUS4, Thermo Fisher). CD10 immunostaining was done on paraffin sections (8 μm) as previously detailed (Perruchot et al, 2016) with the following modifications. After deparaffinization, slides were first incubated with 50mM ammonium chloride (A0171, Sigma-Aldrich) for 10 min and then with 0.1% Sudan black B (s4380, Sigma-Aldrich) in 70% ethanol for 20 min to quench the autofluorescence of immune cells. Slides were then rinsed with Tris-buffered saline (TBS) with 0.02% Tween-20 (P1379, Sigma-Aldrich). Tissue sections were then subjected to heat-induced epitope retrieval in 1mM ethylenediaminetetraacetic acid (EDTA, E9884, Sigma-Aldrich), pH8, using a microwave at 800 watts for 2x5 min. Sections from both frozen and paraffin-embedded tissue were then permeabilized with 0.25% Triton X-100 (T9284, Sigma-Aldrich). Nonspecific-antibody binding was blocked with 2% bovine serum albumin (A2153, Sigma-Aldrich) in TBS. Tissue slices were then sequentially incubated with primary and secondary antibodies (table S1) at 37°C for 1h30 and 45 min, respectively. After washing, nuclei were counterstained with Hoechst 33342 (14533, VWR) at 1 μg/mL for 2 min. Slides were mounted using Vectashield mounting medium (H-1000; Vector Laboratories, Burlingame, CA). Images were obtained with an E400 Nikon microscope (Nikon France, Le Pallet, France) using the NIS-Elements BR4.20.00 software (Nikon).

### Statistical analysis

Data were expressed as means ± SEM. PCR results were subjected to an analysis of variance (ANOVA) using R Studio software. Different letters in Table 2 indicate significant differences (p<0.05 or p<0.01). For statistical analysis of western blot results we used the non-parametric Mann-Whitney *U* test. Significant differences were considered at p<0.05 and trends at p<0.10.

## Acknowledgements

We thank Laurent Deleurme and Gersende Lacombe from the BIOSIT CytomeTRI platform of Rennes (France) for technical assistance. Acknowledgements are also extended to the staff at the INRA dairy farm of Méjusseaume (UMR1348 PEGASE, Le Rheu, France) and Frédérique Mayeur-Nickel for laboratory analyses.

## Author contributions

Laurence Finot performed experiments, data interpretation, statistics and manuscript preparation. Frederic Dessauge supervised project conception, and contributed to the design of experiments and to the writing of the manuscript. Eric Chanat contributed to data interpretation and to the writing of the manuscript.

---

**Fig. S1.**
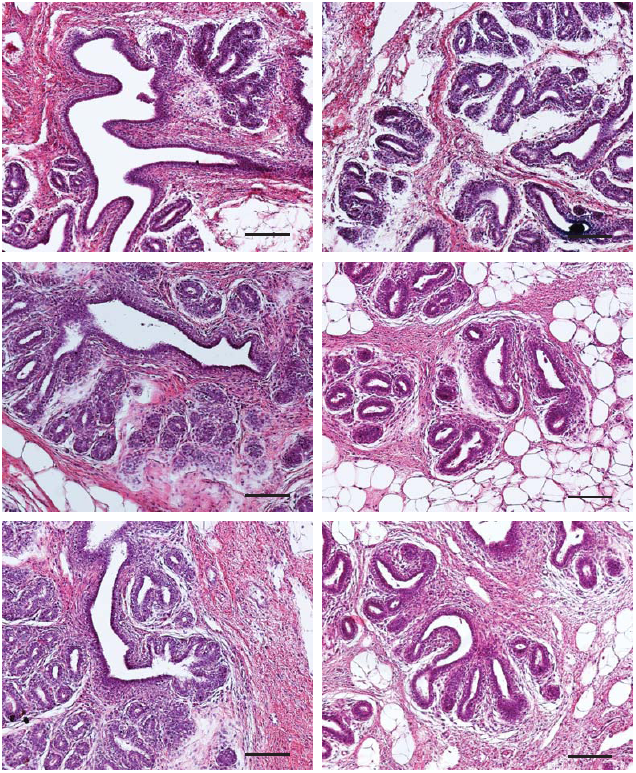
Morphology of bovine mammary tissue at puberty. Mammary tissue fragments from pubertal cows were fixed and processed for histological analysis. Representative tissue sections stained with hematoxylin and eosin are shown. Scale bars: 100 μm.

**Fig. S2.**
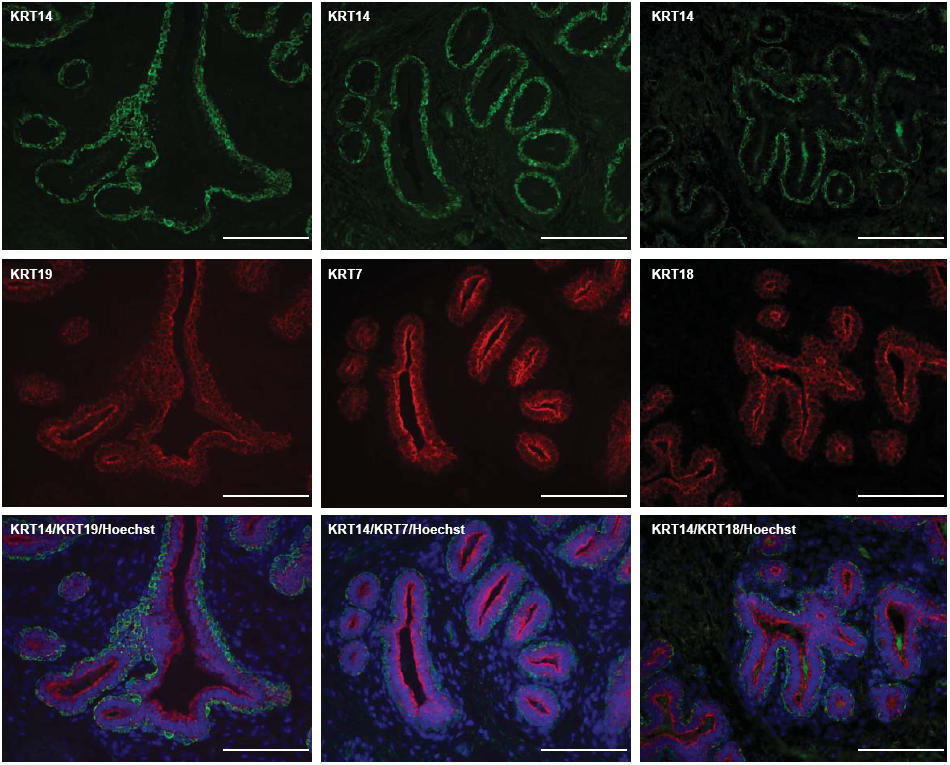
*In situ* localization of keratins demonstrates their lineage-specificity in the developing mammary tissue. Cryo-sections from the mammary tissue of pubertal cows were processed for immunofluorescence for the indicated antigens. Nuclei were counterstained with Hoechst 33342. Keratin 14 (KRT14) was predominantly expressed in basal cells (upper panels, green) whereas KRT19, KRT7 and KRT18 were expressed in luminal cells (middle panels, red). Relative localization of keratins was obtained by image merging of the indicated anti-keratin antibodies (lower panels, color-coded to match the fluorophore). Images are representative of 3 cows. Scale bars, 100μm.

**Fig. S3.**
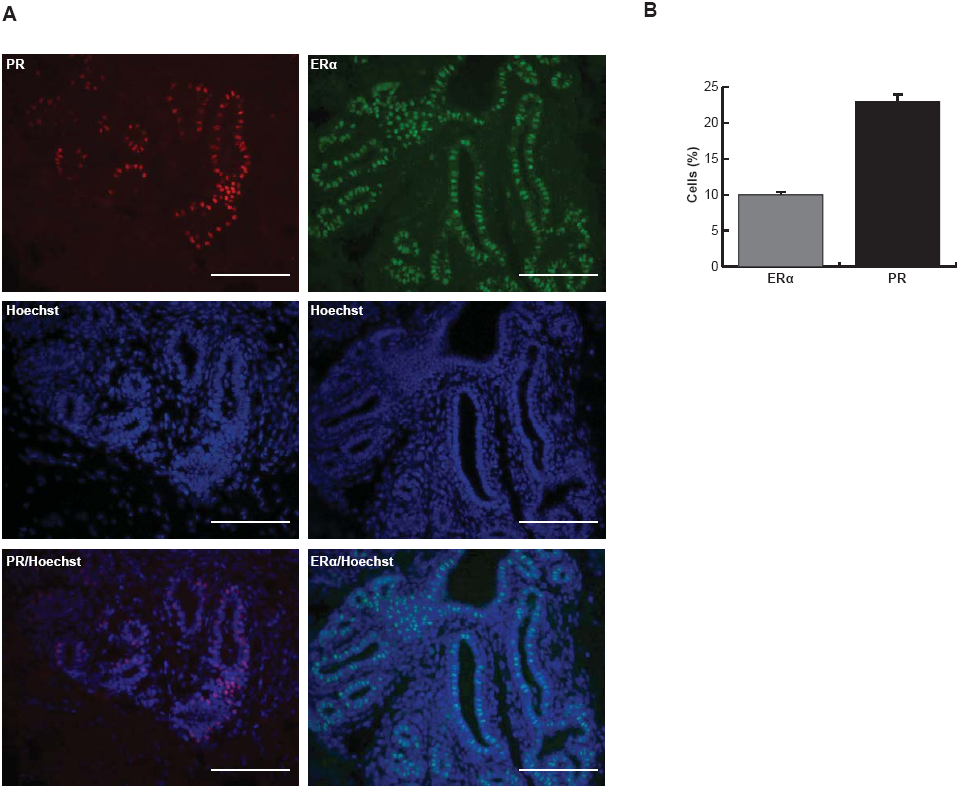
*In situ* localization of the cells expressing the receptors for progesterone and estradiol in the developing mammary tissue. Cryo-sections from the mammary tissue of pubertal cows were processed for immunofluorescence for the progesterone receptor (PR) and estradiol receptor alpha (ERα). Nuclei were counterstained with Hoechst 33342. A) A large number of epithelial cells expressing PR (left panel, red) and ERα (right panel, green) are located in the inner layer of the mammary structures. Images are representative of 3 cows. Scale bars, 100μm. B) Quantification of the cells expressing PR and ERα within the mammary tissue. Results are generated from 6 images per animal for the 3 pubertal cows. Results are given in percentage ±SEM of stained cells (PR or ERα) relative to the total number of cells counterstained with Hoechst 33342.

**Fig. S4.**
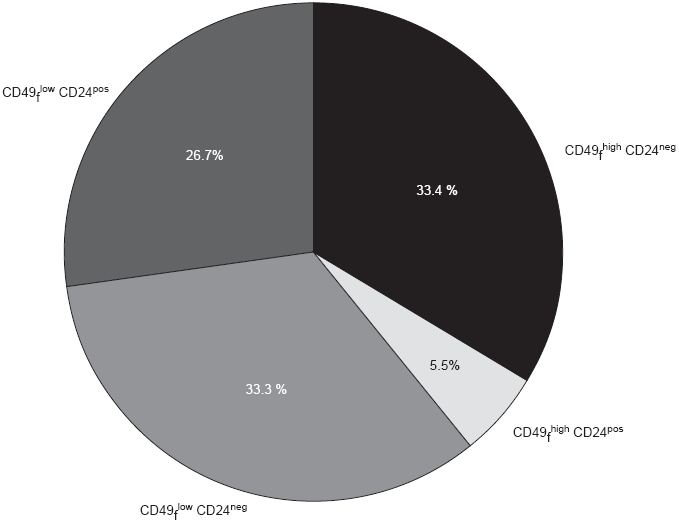
Proportion of each sub-population composing the epithelial cell fraction of the bovine mammary tissue at puberty. Cells dissociated from pubertal bovine mammary tissue were stained with anti-CD49_f_ (CD49_f_) and anti-CD24 (CD24) antibodies, and analyzed by flow cytometry. The number of cells in each sub-population of epithelial cells were expressed as the percentage of the total CD49_f_^low^ orCD49_r_^high^ cells, as shown.

**Table S1.** List of antibodies used for flow cytometry (FACS), Western Blotting and immunofluorescence analyses.

| Antigen | Antibody | Manufacturer | Reference | Dilution (Application) |
| --- | --- | --- | --- | --- |
| CD10 | CD10-PE-Vio770, human (clone 97C5) | Miltenyi Biotec | 130-100-421 | 1:10 (FACS) |
| Isotype control | Mouse IgG1-PE-Vio770 | Miltenyi Biotec | 130-096-654 | 1:10 |
|  | Mouse (clone 56C6) | Dako | M7308 | 1:100 (IF) / 1:2500 (WB) |
| CD24 | CD24-APC, mouse (clone M1/69) | Stem Cell | 60099AZ.1 | 1:10 (FACS) |
| Isotype control | Rat IgG2b-APC | Stem Cell | 60077AZ.1 | 1:10 |
|  | CD24-FITC, mouse (clone M1/69) | Miltenyi Biotec | 130-102-731 | 1:25 (IF) |
| CD49<sub>f</sub> | CD49<sub>f</sub>-FITC, human and mouse (clone GoH3) | Miltenyi Biotec | 130-097-245 | 1:10 (FACS) / 1:25 (IF) |
| Isotype control | Rat IgG2a-FITC | Miltenyi Biotec | 130-102-653 | 1:10 |
|  | CD49<sub>f</sub>-PE, human and mouse (clone GoH3) | Miltenyi Biotec | 130-100-096 | 1:10 (FACS) |
| Isotype control | Rat IgG2a-PE | Miltenyi Biotec | 130-102-654 | 1:10 |
| A-Smooth Muscle (αSMA) | Mouse (clone 1A4) | Santa Cruz | SC32251 | 1:2500 (WB) |
| E-cadherin (CDH1) | Mouse (clone CY-90) | Dako | M3612 | 1:2500 (WB) |
| Estrogen Receptor alpha (ERα) | Rabbit (clone HC-20) | Santa Cruz | SC543 | 1:100 (IF) |
| Keratin 7 (KRT7) | Mouse (clone 5F282) | Santa Cruz | SC70936 | 1:2500 (WB) |
| Keratin 14 (KRT14) | Goat (clone C-14) | Santa Cruz | SC17104 | 1:2500 (WB) |
| Keratin 18 (KRT18) | Mouse (clone NCH38) | Sigma-Aldrich | C8541-.2ML | 1:2500 (WB) |
| Keratin 19 (KRT19) | Mouse (clone b170) | Leica Biosystems | NCL-CK19 | 1:2500 (WB) |
| Progesterone Receptor (PR) | Mouse (clone PR10A9) | Beckman Coulter | PN IM1546 | 1:200 (IF) |

**Table S2.**
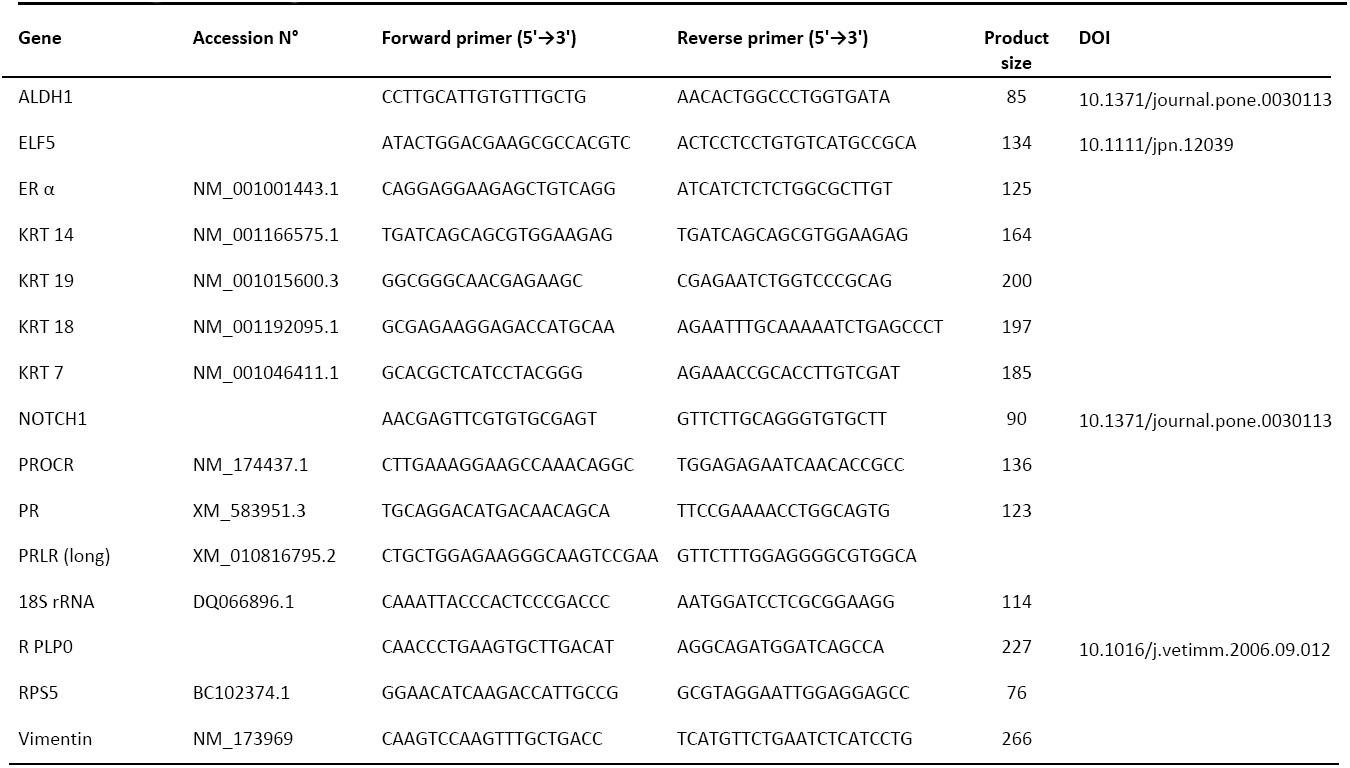
List of primers used in quantitative PCR

## REFERENCES

Akers, R. M. (2017) ‘A 100-Year Review: Mammary development and lactation’, Journal of dairy science 100(12): 10332–10352.

Arevalo Turrubiarte, M., Perruchot, M. H., Finot, L., Mayeur, F. and Dessauge, F. (2016) ‘Phenotypic and functional characterization of two bovine mammary epithelial cell lines in 2D and 3D models’, American journal of physiology. Cell physiology 310(5): C348–56.

Asselin-Labat, M. L., Shackleton, M., Stingl, J., Vaillant, F., Forrest, N. C., Eaves, C. J., Visvader, J. E. and Lindeman, G. J. (2006) ‘Steroid hormone receptor status of mouse mammary stem cells’, Journal of the National Cancer Institute 98(14): 1011–4.

Asselin-Labat, M. L., Vaillant, F., Shackleton, M., Bouras, T., Lindeman, G. J. and Visvader, J. E. (2008) ‘Delineating the epithelial hierarchy in the mouse mammary gland’, Cold Spring Harbor symposia on quantitative biology 73: 469–78.

Atwood, C. S., Hovey, R. C., Glover, J. P., Chepko, G., Ginsburg, E., Robison, W. G. and Vonderhaar, B. K. (2000) ‘Progesterone induces side-branching of the ductal epithelium in the mammary glands of peripubertal mice’, The Journal of endocrinology 167(1): 39–52.

Bachelard-Cascales, E., Chapellier, M., Delay, E., Pochon, G., Voeltzel, T., Puisieux, A., Caron de Fromentel, C. and Maguer-Satta, V. (2010) ‘The CD10 enzyme is a key player to identify and regulate human mammary stem cells’, Stem cells 28(6): 1081–8.

Borena, B. M., Bussche, L., Burvenich, C., Duchateau, L. and Van de Walle, G. R. (2013) ‘Mammary stem cell research in veterinary science: an update’, Stem cells and development 22(12): 1743–51.

Brisken, C. and Ataca, D. (2015) ‘Endocrine hormones and local signals during the development of the mouse mammary gland’, Wiley interdisciplinary reviews. Developmental biology 4(3): 181–95.

Capuco, A. V., Choudhary, R. K., Daniels, K. M., Li, R. W. and Evock-Clover, C. M. (2012) ‘Bovine mammary stem cells: cell biology meets production agriculture’, Animal : an international journal of animal bioscience 6(3): 382–93.

Chakrabarti, R., Wei, Y., Romano, R. A., DeCoste, C., Kang, Y. and Sinha, S. (2012) ‘Elf5 regulates mammary gland stem/progenitor cell fate by influencing notch signaling’, Stem cells 30(7): 1496–508.

Dairkee, S. H., Puett, L. and Hackett, A. J. (1988) ‘Expression of basal and luminal epithelium-specific keratins in normal, benign, and malignant breast tissue’, Journal of the National Cancer Institute 80(9): 691–5.

Deome, K. B., Faulkin, L. J., Jr., Bern, H. A. and Blair, P. B. (1959) ‘Development of mammary tumors from hyperplastic alveolar nodules transplanted into gland-free mammary fat pads of female C3H mice’, Cancer research 19(5): 515–20.

Dontu, G. and Ince, T. A. (2015) ‘Of mice and women: a comparative tissue biology perspective of breast stem cells and differentiation’, Journal of mammary gland biology and neoplasia 20(1–2): 51–62.

Eirew, P., Kannan, N., Knapp, D. J., Vaillant, F., Emerman, J. T., Lindeman, G. J., Visvader, J. E. and Eaves, C. J. (2012) ‘Aldehyde dehydrogenase activity is a biomarker of primitive normal human mammary luminal cells’, Stem cells 30(2): 344–8.

Hilton, H. N., Graham, J. D., Kantimm, S., Santucci, N., Cloosterman, D., Huschtscha, L. I., Mote, P. A. and Clarke, C. L. (2012) ‘Progesterone and estrogen receptors segregate into different cell subpopulations in the normal human breast’, Molecular and cellular endocrinology 361(1–2): 191–201.

Inman, J. L., Robertson, C., Mott, J. D. and Bissell, M. J. (2015) ‘Mammary gland development: cell fate specification, stem cells and the microenvironment’, Development 142(6): 1028–42.

Lichtner, R. B., Julian, J. A., North, S. M., Glasser, S. R. and Nicolson, G. L. (1991) ‘Coexpression of cytokeratins characteristic for myoepithelial and luminal cell lineages in rat 13762NF mammary adenocarcinoma tumors and their spontaneous metastases’, Cancer research 51(21): 5943–50.

Martignani, E., Eirew, P., Accornero, P., Eaves, C. J. and Baratta, M. (2010) ‘Human milk protein production in xenografts of genetically engineered bovine mammary epithelial stem cells’, PloS one 5(10): e13372.

Martignani, E., Eirew, P., Eaves, C. and Baratta, M. (2009) ‘Functional identification of bovine mammary epithelial stem/progenitor cells’, Veterinary research communications 33 Suppl 1: 101–3.

McBryan, J. and Howlin, J. (2017) ‘Pubertal Mammary Gland Development: Elucidation of In Vivo Morphogenesis Using Murine Models’, Methods in molecular biology 1501: 77–114.

O’Leary, K. A., Shea, M. P., Salituro, S., Blohm, C. E. and Schuler, L. A. (2017) ‘Prolactin Alters the Mammary Epithelial Hierarchy, Increasing Progenitors and Facilitating Ovarian Steroid Action’, Stem cell reports 9(4): 1167–1179.

Oakes, S. R., Naylor, M. J., Asselin-Labat, M. L., Blazek, K. D., Gardiner-Garden, M., Hilton, H. N., Kazlauskas, M., Pritchard, M. A., Chodosh, L. A., Pfeffer, P. L. et al. (2008) ‘The Ets transcription factor Elf5 specifies mammary alveolar cell fate’, Genes & development 22(5): 581–6.

Ormandy, C. J., Naylor, M., Harris, J., Robertson, F., Horseman, N. D., Lindeman, G. J., Visvader, J. and Kelly, P. A. (2003) ‘Investigation of the transcriptional changes underlying functional defects in the mammary glands of prolactin receptor knockout mice’, Recent progress in hormone research 58: 297–323.

Ormerod, E. J. and Rudland, P. S. (1986) ‘Regeneration of mammary glands in vivo from isolated mammary ducts’, Journal of embryology and experimental morphology 96: 229–43.

Osinska, E., Wicik, Z., Godlewski, M. M., Pawlowski, K., Majewska, A., Mucha, J., Gajewska, M. and Motyl, T. (2014) ‘Comparison of stem/progenitor cell number and transcriptomic profile in the mammary tissue of dairy and beef breed heifers’, Journal of applied genetics 55(3): 383–95.

Perruchot, M. H., Arevalo-Turrubiarte, M., Dufreneix, F., Finot, L., Lollivier, V., Chanat, E., Mayeur, F. and Dessauge, F. (2016) ‘Mammary Epithelial Cell Hierarchy in the Dairy Cow Throughout Lactation’, Stem cells and development 25(19): 1407–18.

Peuhu, E., Virtakoivu, R., Mai, A., Warri, A. and Ivaska, J. (2017) ‘Epithelial vimentin plays a functional role in mammary gland development’, Development.

Prater, M. D., Petit, V., Alasdair Russell, I., Giraddi, R. R., Shehata, M., Menon, S., Schulte, R., Kalajzic, I., Rath, N., Olson, M. F. et al. (2014) ‘Mammary stem cells have myoepithelial cell properties’, Nature cell biology 16(10): 942–50, 1–7.

Rauner, G. and Barash, I. (2012) ‘Cell hierarchy and lineage commitment in the bovine mammary gland’, PloS one 7(1): e30113.

Rauner, G. and Barash, I. (2016) ‘Enrichment for Repopulating Cells and Identification of Differentiation Markers in the Bovine Mammary Gland’, Journal of mammary gland biology and neoplasia 21(1–2): 41–9.

Rauner, G., Ledet, M. M. and Van de Walle, G. R. (2017) ‘Conserved and variable: Understanding mammary stem cells across species’, Cytometry. Part A : the journal of the International Society for Analytical Cytology.

Safayi, S., Korn, N., Bertram, A., Akers, R. M., Capuco, A. V., Pratt, S. L. and Ellis, S. (2012) ‘Myoepithelial cell differentiation markers in prepubertal bovine mammary gland: effect of ovariectomy’, Journal of dairy science 95(6): 2965–76.

Scheele, C. L., Hannezo, E., Muraro, M. J., Zomer, A., Langedijk, N. S., van Oudenaarden, A., Simons, B. D. and van Rheenen, J. (2017) ‘Identity and dynamics of mammary stem cells during branching morphogenesis’, Nature 542(7641): 313–317.

Seagroves, T. N., Lydon, J. P., Hovey, R. C., Vonderhaar, B. K. and Rosen, J. M. (2000) ‘C/EBPbeta (CCAAT/enhancer binding protein) controls cell fate determination during mammary gland development’, Molecular endocrinology 14(3): 359–68.

Shackleton, M., Vaillant, F., Simpson, K. J., Stingl, J., Smyth, G. K., Asselin-Labat, M. L., Wu, L., Lindeman, G. J. and Visvader, J. E. (2006) ‘Generation of a functional mammary gland from a single stem cell’, Nature 439(7072): 84–8.

Shehata, M., Teschendorff, A., Sharp, G., Novcic, N., Russell, I. A., Avril, S., Prater, M., Eirew, P., Caldas, C., Watson, C. J. et al. (2012) ‘Phenotypic and functional characterisation of the luminal cell hierarchy of the mammary gland’, Breast cancer research : BCR 14(5): R134.

Shipitsin, M., Campbell, L. L., Argani, P., Weremowicz, S., Bloushtain-Qimron, N., Yao, J., Nikolskaya, T., Serebryiskaya, T., Beroukhim, R., Hu, M. et al. (2007) ‘Molecular definition of breast tumor heterogeneity’, Cancer Cell 11(3): 259–73.

Sleeman, K. E., Kendrick, H., Ashworth, A., Isacke, C. M. and Smalley, M. J. (2006) ‘CD24 staining of mouse mammary gland cells defines luminal epithelial, myoepithelial/basal and non-epithelial cells’, Breast cancer research : BCR 8(1): R7.

Smith, G. H. and Medina, D. (1988) ‘A morphologically distinct candidate for an epithelial stem cell in mouse mammary gland’, Journal of cell science 90 (Pt 1): 173–83.

Stingl, J. (2009) ‘Detection and analysis of mammary gland stem cells’, The Journal of pathology 217(2): 229–41.

Stingl, J., Eirew, P., Ricketson, I., Shackleton, M., Vaillant, F., Choi, D., Li, H. I. and Eaves, C. J. (2006) ‘Purification and unique properties of mammary epithelial stem cells’, Nature 439(7079): 993–7.

Van Keymeulen, A., Rocha, A. S., Ousset, M., Beck, B., Bouvencourt, G., Rock, J., Sharma, N., Dekoninck, S. and Blanpain, C. (2011) ‘Distinct stem cells contribute to mammary gland development and maintenance’, Nature 479(7372): 189–93.

Villadsen, R., Fridriksdottir, A. J., Ronnov-Jessen, L., Gudjonsson, T., Rank, F., LaBarge, M. A., Bissell, M. J. and Petersen, O. W. (2007) ‘Evidence for a stem cell hierarchy in the adult human breast’, The Journal of cell biology 177(1): 87–101.

Visvader, J. E. and Stingl, J. (2014) ‘Mammary stem cells and the differentiation hierarchy: current status and perspectives’, Genes & development 28(11): 1143–58.

Wang, D., Cai, C., Dong, X., Yu, Q. C., Zhang, X. O., Yang, L. and Zeng, Y. A. (2015) ‘Identification of multipotent mammary stem cells by protein C receptor expression’, Nature 517(7532): 81–4.

Yart, L., Lollivier, V., Marnet, P. G. and Dessauge, F. (2014) ‘Role of ovarian secretions in mammary gland development and function in ruminants’, Animal : an international journal of animal bioscience 8(1): 72–85.

